# A sequence-based approach for identifying protein fold switchers

**DOI:** 10.1101/462606

**Authors:** Soumya Mishra, Loren L. Looger, Lauren L. Porter

**Author notes:** To whom correspondence should be addressed: Lauren Porter, Howard Hughes Medical Institute, Janelia Research Campus, 19700 Helix Dr. Ashburn, VA 20147.

## Abstract

Although most proteins conform to the classical one-structure/one-function paradigm, an increasing number of proteins with dual structures and functions are emerging. These fold-switching proteins remodel their secondary structures in response to cellular stimuli, fostering multi-functionality and tight cellular control. Accurate predictions of fold-switching proteins could both suggest underlying mechanisms for uncharacterized biological processes and reveal potential drug targets. Previously, we developed a prediction method for fold-switching proteins based on secondary structure predictions and structure-based thermodynamic calculations. Given the large number of genomic sequences without homologous experimentally characterized structures, however, we sought to predict fold-switching proteins from their sequences alone. To do this, we leveraged state-of-the-art secondary structure predictions, which require only amino acid sequences but are not currently designed to identify structural duality in proteins. Thus, we hypothesized that incorrect and inconsistent secondary structure predictions could be good initial predictors of fold-switching proteins. We found that secondary structure predictions of fold-switching proteins with solved structures are indeed less accurate than secondary structure predictions of non-fold-switching proteins with solved structures. These inaccuracies result largely from the conformations of fold-switching proteins that are underrepresented in the Protein Data Bank (PDB), and, consequently, the training sets of secondary structure predictors. Given that secondary structure predictions are homology-based, we hypothesized that decontextualizing the inaccurately-predicted regions of fold-switching proteins could weaken the homology relationships between these regions and their overpopulated structural representatives. Thus, we reran secondary structure predictions on these regions in isolation and found that they were significantly more inconsistent than in regions of non-fold-switching proteins. Thus, inconsistent secondary structure predictions can serve as a preliminary marker of fold switching. These findings have implications for genomics and the future development of secondary structure predictors.

## Introduction

Most structurally characterized proteins perform one well-defined function supported by one scaffold of secondary structure [1]. (Proteins are highly dynamic molecules[2] at the tertiary level, but typically not at the secondary level.) This observation, substantiated by over 100,000 atomic-level protein structures [3], has laid the basis for homology-based protein structure prediction algorithms. These algorithms are powerful enough to predict the tertiary structures of previously uncharacterized proteins with significant sequence similarity to a solved protein structure. If the sequence of a protein of interest falls below the threshold of significant similarity [4], sequence-based secondary structure predictions remain a viable option for model building. These lower-resolution predictions can be combined with other methods to develop putative three-dimensional protein structures [5-7], and can even suggest their possible functional repertoire [8, 9].

Recent work has shown, however, that many proteins substantially remodel their secondary structures in response to cellular stimuli, enabling radical functional changes and tight cellular control [10]. This phenomenon, called fold switching [11], can involve structural and functional transformations as drastic as an α-helical transcription factor morphing into a β-barrel translation factor [12]. Additionally, some fold-switching proteins change their cellular localization. For example, Chloride intracellular channel protein 1 (CLIC1) is a human protein that functions as both a cytosolic glutathione reductase [13] and a membrane-inserted chloride channel [14]. While entire protein domains can switch folds [12], current experimental evidence suggests that it is more common for subdomains of larger proteins to switch folds while the remainder maintains a single native structure. We call the structurally changing subdomains “fold-switching regions” (FSRs) and the structurally unchanging remainders “non-fold-switching regions” (NFSRs).

Predicting the fold-switching ability of a given protein region can suggest a mechanism for its function *in situ*, especially when combined with other forms of evidence that the protein is multifunctional, has more than one cellular localization, *etc*. However, it is not yet possible to correctly make such predictions with confidence. Many factors contribute to this situation. All available secondary structure prediction methods return a single prediction for a given amino acid sequence, without even an estimate of the confidence of the prediction. Homology-based tertiary predictions methods are currently unable to predict multiple distinct conformations of a sequence. Tertiary structure prediction tools that incorporate *de novo* folding elements, such as Rosetta[15, 16], can return an ensemble of 3-dimensional models. Such ensembles can be viewed either as multiple guesses at the correct structure or as an estimate of the dynamic conformational rearrangements around a core prediction. However, even these state-of-the-art algorithms are unable to deal with multiple potentially well-folded backbone scaffolds [17], and thus cannot adequately predict fold switching either.

Previously, we successfully predicted FSRs by exploiting the failures of homology-based secondary structure predictors [10]. Specifically, we showed that discrepancies between predicted and experimentally determined secondary structures can indicate that a given protein switches folds. By coupling these discrepancies with a structure-based thermodynamic calculation [18], we were able to successfully predict fold switching in 13 proteins with one solved structure and experimental evidence for an alternative conformation. Thus, discrepancies between predicted and experimentally determined secondary structures can contribute (along with thermodynamic calculations and literature evidence) to estimating whether a given amino acid sequence switches folds.

Here, we seek to determine if secondary structure discrepancies alone can productively suggest whether or not a given protein switches folds. This approach has the advantage of not requiring a solved protein structure with significant sequence similarity to the protein of interest. To lay the basis for this approach, we show that incorrect secondary structure predictions are significantly more common within FSRs than within NFSRs. We find that these incorrect predictions arise predominantly from one of the two conformations accessible to a given fold-switching protein. Inaccurately predicted conformers tend to be underrepresented in the Protein Data Bank (PDB), demonstrating that secondary structure predictions are influenced by structural bias within the PDB. We show that this bias can be leveraged to identify FSRs using secondary structure prediction discrepancies between FSR sequences in isolation and identical sequences within the context of their parent proteins. Thus, secondary structure discrepancies can be used as a preliminary predictor of fold switching from amino acid sequence alone. This result has implications for the improvement of secondary structure predictors as well as identification of fold switching in proteins of biological interest.

## Results

### Secondary structure predictions of fold-switching proteins are consistently less accurate than those of non-fold switchers

First, we compared secondary structure prediction accuracies of FSRs with those of NFSRs. To do this, we ran three secondary structure prediction software packages (JPred [19], PSIPred [20], and SPIDER2 [21]) on a curated set of 192 structures of fold-switching proteins [10] (96 proteins, 2 structures each) and compared the predicted secondary structure annotations of FSRs and NFSRs with the corresponding experimentally determined secondary structures. **Fig. 1a-c** shows the prediction accuracy distributions of both fold-switching and non-fold-switching protein regions. For all three algorithms, the predictions of fold-switching regions were significantly less accurate than the predictions of non-fold-switching regions, with p-values of 8.6*10^−309^, 2.2*10^−179^, and 7.1*10^−135^, respectively (Kolmogorov-Smirnov Test). Average prediction accuracy for FSRs was 67-68%, whereas accuracy for NFSRs was 74-80% (**Table 1**).

**Figure 1.**
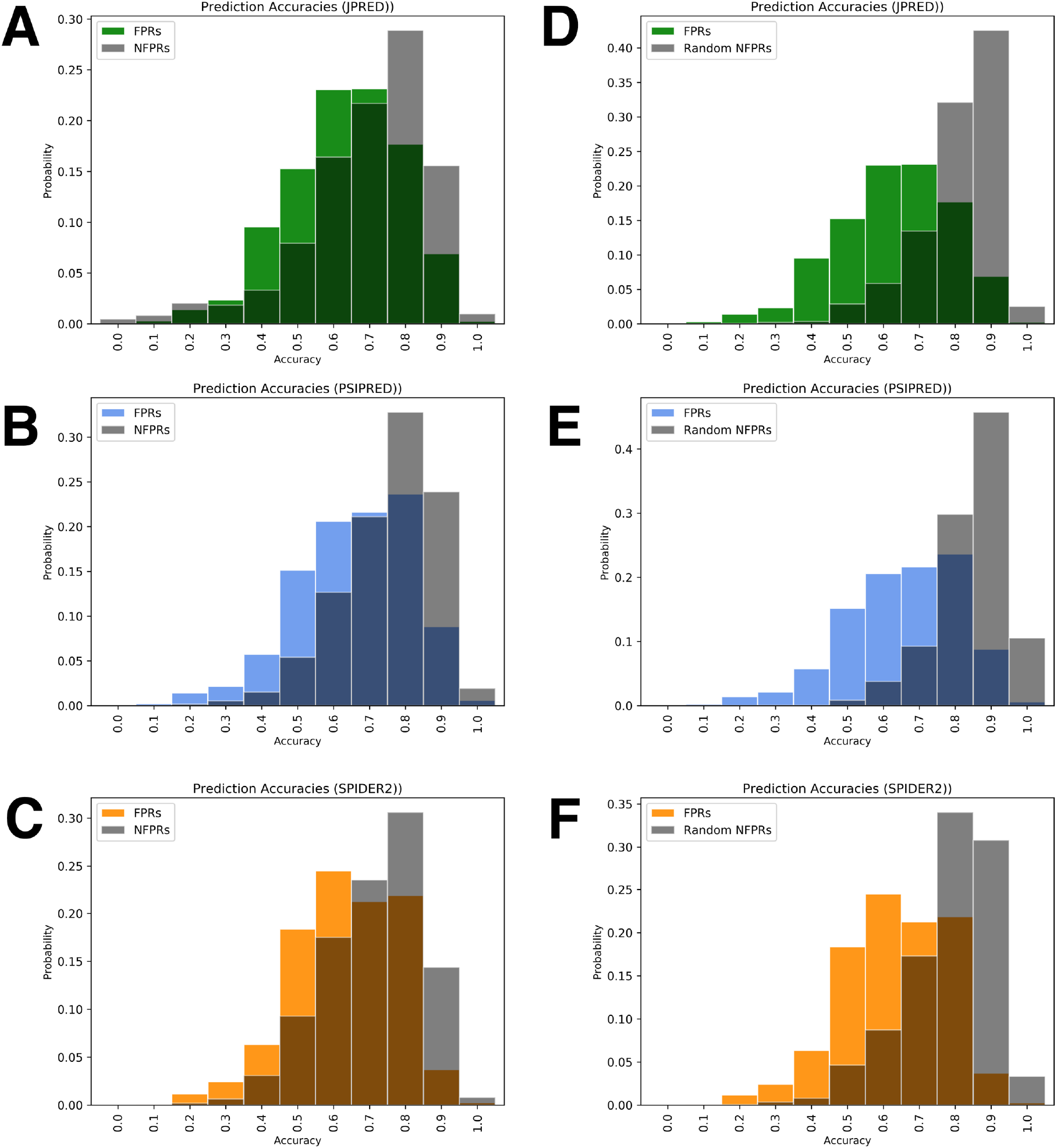
Secondary structure predictions of FSRs are consistently less accurate than those of NFSRs (a-c) and randomly-selected NFSRs (d-f). Histograms of fold-switching fragments are colored (green, JPRED; blue, PSIPRED; orange, SPIDER2), while comparisons of non-fold-switching fragments from corresponding predictors are gray.

**Table 1.** Mean and median secondary structure prediction accuracies

|  | JPred mean/median | PSIPRED mean/median | SPIDER2 mean/median |
| --- | --- | --- | --- |
| FSRs | 0.67/0.69 | 0.68/0.71 | 0.67/0.68 |
| NFSRs | 0.74/0.79 | 0.80/0.83 | 0.76/0.78 |
| Random NFSRs | 0.85/0.88 | 0.89/0.9 | 0.83/0.85 |
| Better FSRs | 0.76/0.77 | 0.75/0.77 | 0.72/0.74 |

Next, we randomly selected regions of proteins that are expected not to switch folds [10] and calculated secondary structure prediction accuracy using their solved structures (**Fig. 1 d-f**). The secondary structures of these fragments were also predicted much more accurately than for FSRs, with p-values of 1.2*10^−264^, 1.6*10^−276^, and 3.3*10^−176^, respectively (Kolmogorov-Smirnov Test). Average prediction accuracy for the random NFSRs was 83-89% (**Table 1**), even better than that for the NFSRs from our curated fold-switching proteins. Together, these results indicate that secondary structure prediction accuracies of FSRs are significantly worse than the corresponding accuracies of NFSRs, using three state-of-the-art prediction algorithms. This was true both for static regions of fold-switching proteins and for randomly selected regions of non-fold-switching proteins.

### Secondary structure prediction inaccuracies in FSRs result largely from one conformation

It is perhaps unsurprising that secondary structure predictions of FSRs are significantly less accurate than NFSRs. After all, JPRED, PSIPRED, and SPIDER2 are designed to predict a single, best-guess secondary structure configuration for a given amino acid sequence. Since by definition FSRs have two distinct secondary structure configurations, single-prediction algorithms could at best correctly predict half of the FSR configurations. Alternatively, they could predict the two FSR configurations equally inaccurately.

Instead, we hypothesized that the algorithms might produce predictions for a given FSR sequence that were significantly closer to one of the two experimentally determined configurations than the other. To test this, we remade our secondary structure prediction accuracy distributions for FSRs by including only the more accurate prediction of the two conformations (**Fig. 2**). We found that the resulting FSR accuracy distributions were significantly more similar to those of NFSRs than to all FSRs (**Table 1**), with p-values indicating similar distributions within the realm of chance (0.88, 0.11, and 0.08, respectively, Kolmogorov-Smirnov test). Thus, JPRED, PSIPRED, and SPIDER2 tend to predict one FSR conformation with reasonable accuracy but predict the alternative poorly. The remaining discrepancies between FSR and NFSR inaccuracies result from SS predictor tendencies to predict the secondary structures of both FSR conformations inaccurately. This accounts for 27%, 32% and 40% of the predictions from JPRED, PSIPRED, and SPIDER2, respectively.

**Figure 2.**
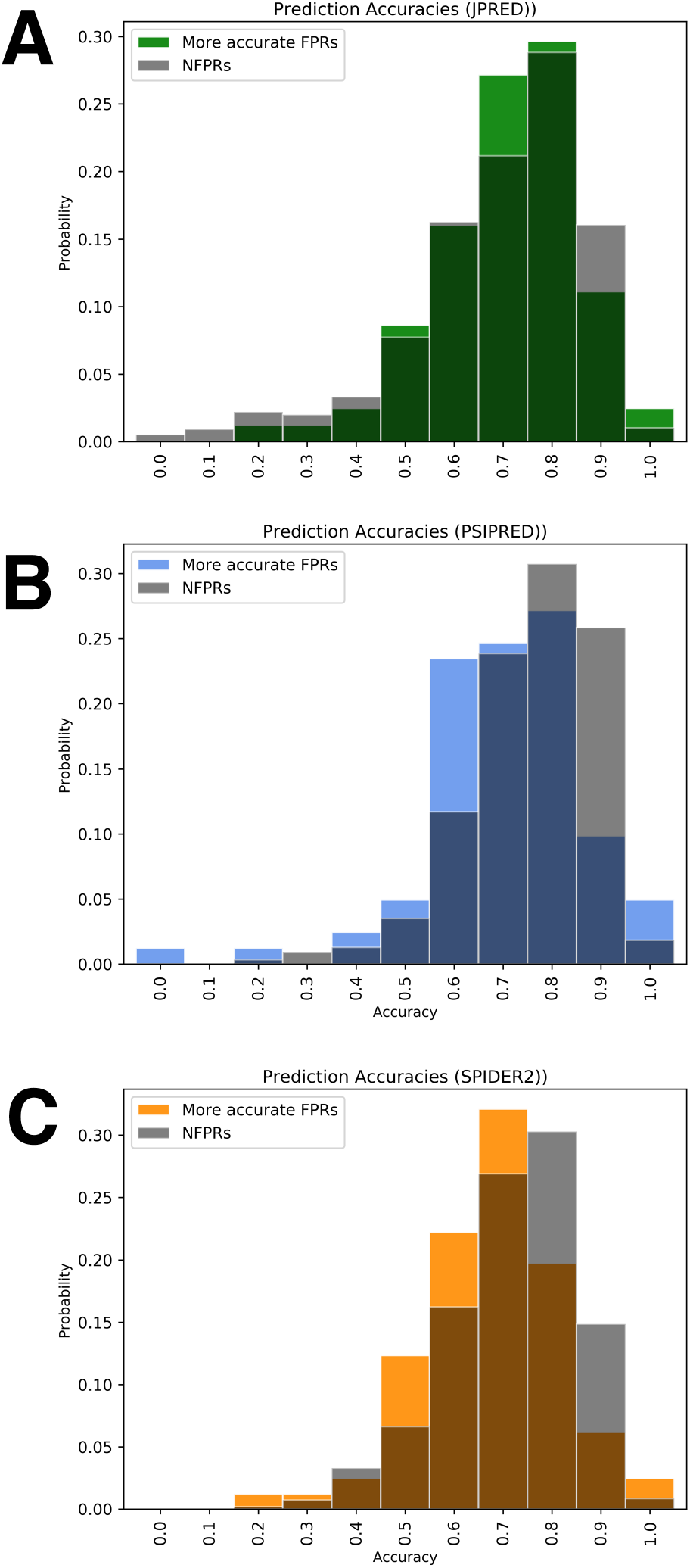
Better FSR predictions are significantly more accurate and similar to NFSR distributions. Histograms of fold-switching fragments are colored (green, JPRED; blue, PSIPRED; orange, SPIDER2), while comparisons of non-fold-switching fragments from corresponding predictors are gray.

### Conformational overrepresentation contributes to incorrect secondary structure predictions of FSRs

Upon noticing FSR secondary structure prediction bias (*i.e.* predictions looking significantly more like one conformation than the other), we sought to determine its source. We hypothesized that FSRs with high frequencies within the PDB were more likely to be predicted correctly, while FSRs with fewer representative structures were more likely to be predicted incorrectly. This hypothesis was based on the fact that these three secondary structure predictors are trained on proteins with previously solved structures. Thus, training sets would more likely contain protein conformations that were highly represented within the PDB. Alternatively, the observed prediction bias might result from differences in the algorithms’ performance on the distinct secondary structures, for instance.

To test our hypothesis that asymmetric FSR occurrence within the PDB leads secondary structure predictors to predict one FSR conformation more accurately than the other, we determined the number of structures available for each FSR conformation (**Supp. Table 1**) and tested if FSRs with more accurate predictions tended to have more representative structures in the PDB. Our null hypothesis was that FSRs were equally likely to be predicted correctly, regardless of how many representative structures they had. By performing the Binomial Test, we disproved the null hypothesis (p-values of 7.7*10^−4^, 8.2*10^−3^, and 6.3*10^−2^, respectively), demonstrating that secondary structure predictions are biased towards predicting FSR conformations with frequent PDB representation and biased against predicting FSR conformations with infrequent PDB representation.

### Secondary structure predictions of FSRs are context-dependent

Given that the secondary structure predictions of these three algorithms depend heavily on homology-based sequence inferences [22], we hypothesized that removing the sequence context from around fold-switching protein regions could result in alternative secondary structure predictions. This hypothesis was based computational and experimental evidence indicating that secondary structure formation can be context-dependent [23-25]. Thus, we ran secondary structure predictions on the amino acid sequences encoding FSRs without the flanking NFSR sequences. As expected, predictions of isolated FSR fragments differed from predictions of FSRs in the context of the whole protein. However, the overall distributions these discerpancies were similar to those of randomly-selected fragments of equivalent length from non-switching proteins. Distributions differed upon regeneration using only FSR fragments whose in-context prediction accuracies were < 0.8, the approximate average accuracy for all three secondary structure predictors [19, 21, 26] (**Fig. 3a-c, Table 2**, statistical significances of 1.3*10^−4^ (JPRED), 2.2*10-2 (PSIPRED), and 3.7*10^−7^ (SPIDER2)). These results indicate that discrepancies in secondary structure predictions of protein fragments in isolation *vs*. the context of the whole protein are a good initial predictor of fold switching if secondary structure predictions for both conformers are not especially accurate.

**Figure 3.**
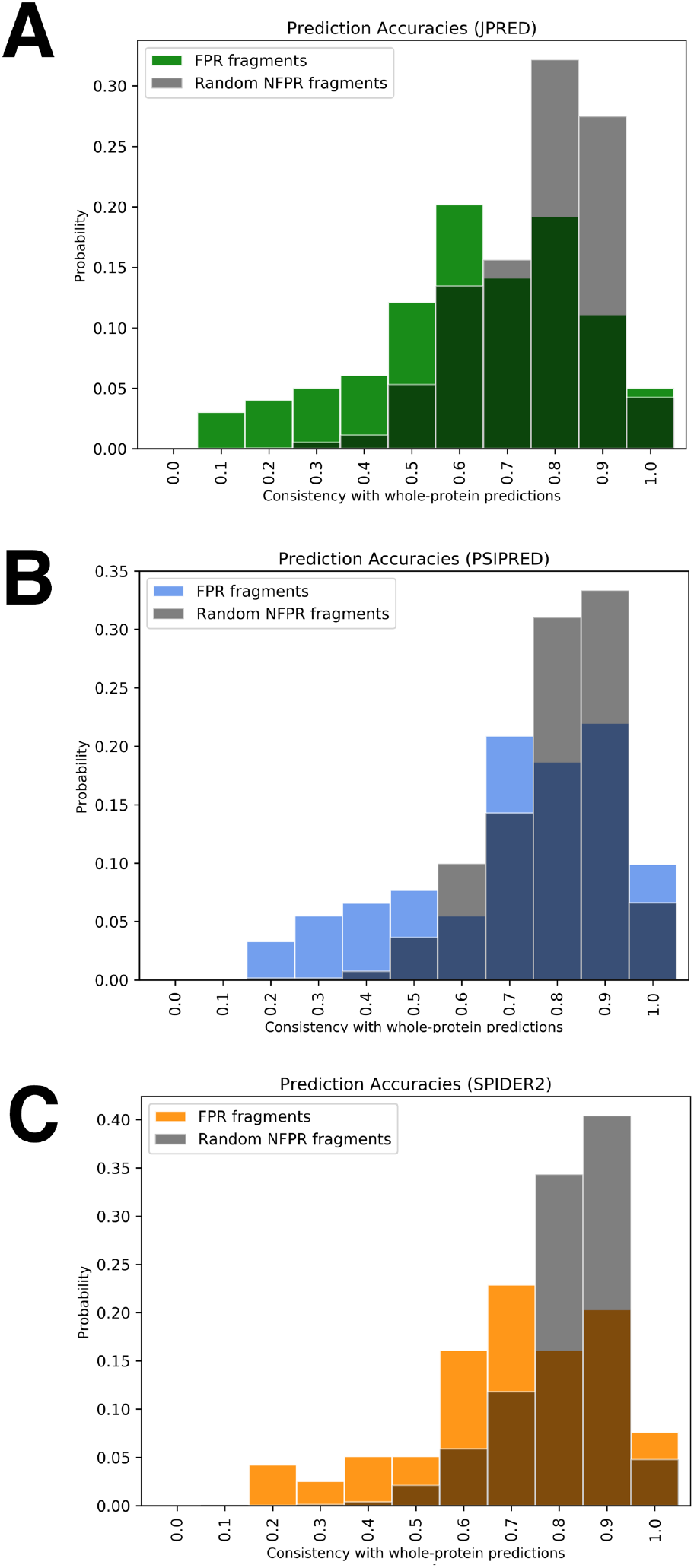
Consistencies between secondary structures of protein fragments in isolation and in the context of the whole protein. Comparisons of fold-switching fragments are colored (green, JPRED; blue, PSIPRED; orange, SPIDER2), while comparisons of non-fold-switching fragments from corresponding predictors are gray.

**Table 2.** Mean and median secondary structure prediction comparisons (fragment:whole)

|  | JPred mean/median | PSIPRED mean/median | SPIDER2 mean/median |
| --- | --- | --- | --- |
| FSRs ($PA^1 < 0.8$, both conformers) | 0.68/0.7 | 0.75/0.80 | 0.75/0.77 |
| NFSRs | 0.81/0.83 | 0.84/0.85 | 0.86/0.88 |

<sup>1</sup>PA = secondary structure Prediction Accuracies

## Discussion

While most proteins with solved structures adhere to the classical notion that proteins adopt one secondary structure scaffold that performs one specific function, there are a number of exceptions [10, 27]. Such exceptions include fold-switching proteins, which remodel their secondary structures in response to cellular stimuli, fostering changes in function or enabling tight cellular control. Because fold-switching proteins do not conform to the classical notion, their dual conformations and functionalities are unlikely to be recognized by current homology-based secondary structure predictors. Here, we seek to leverage this observation by using faulty secondary structure predictions to identify fold-switching proteins. Similar calculations that collate secondary structure predictions to identify flexible regions in proteins have been performed previously, though on a very limited dataset [28].

Here we show that inaccurate and inconsistent secondary structure predictions can reveal fold-switching protein regions. We first showed that secondary structure predictions that are inconsistent with experimentally determined protein structures are significantly more common in FSRs than in non-FSRs. This observation gives statistical power to our previous observation that secondary structure predictions of FSRs are often inconsistent with experiment. We then suggested a new predictive approach by showing that secondary structure predictions of FSRs are often context-dependent. That is, secondary structure predictions of FSRs in isolation often differ from those in context. These context-dependent differences occur significantly more frequently in FSRs than in NFSRs. Thus, these differences can be used as a preliminary indicator of fold switching that relies on amino acid sequence alone, as opposed to previous calculations[10], which also require one solved structure from a homolog.

In retrospect, it is not surprising that secondary structure predictions of FSRs are context-dependent. A number of computational and experimental studies have highlighted that structural context can determine secondary structure. Pioneering work by Minor and Kim demonstrated that a “chameleon sequence” of eight amino acids could fold into an α-helix in one part of protein G, but a β-hairpin in another [23]. Furthermore, several computational studies have scoured the PDB to find dozens of naturally-occurring chameleon sequences, with an upper limit of 20 residues [29-31]. Finally, a recent study leveraged context dependence to design 56-amino-acid sequences with 80% sequence identity but different folds [24]. Such chameleon sequences and “near-chameleon sequences” will likely confound homology-based prediction methods, particularly of small fragments.

Although context-dependent secondary structure predictions are a good preliminary indicator of fold switching, they are not definitive: some NFSRs also show discrepancies between isolated and in-context predictions. Other factors—such as independent folding cooperativity—appear necessary for proteins to switch folds [10]. Furthermore, current secondary structure prediction algorithms depend heavily on available amino acid sequences of proteins with solved structures. Our results suggest that this dependence can lead to prediction bias when one configuration of a fold-switching protein is overrepresented relative to another. Protein structure predictions based on physical principles [32] —instead of homology relationships—could circumvent this bias. Thus, we are optimistic that as physically-based secondary structure predictions improve, fold-switching proteins will become easier to predict.

Accurate predictions of fold switching could suggest biological mechanisms underlying observed experimental phenomena. For example, some proteins can change their cellular localizations by switching folds [14, 33]. Others require fold switching to change their functions [12, 34]. Thus, predictions suggesting that a protein switches folds could lead to the identification of mechanisms underlying currently unexplained biological processes. Furthermore, fold-switching proteins could constitute promising drug targets if their conformational equilibria could be disrupted by small molecules. Consistent with this notion, the veterinary medicine halofuginone arrests growth of the malaria parasite through the stabilization of a prolyl tRNA synthetase in an inactive configuration [35, 36]. Thus, predicting whether a protein switches folds could foster the discovery of new biological processes and drug discovery paradigms. We hope that the results presented here will help to lay the groundwork for these advances.

## Methods

### Secondary structure predictions of whole fold-switching proteins

All full amino acid sequences of 192 fold-switching protein structures [10], corresponding to two different conformations of 96 fold-switching proteins (96 proteins, 2 structures each; aka fold-switch pairs), were downloaded from the PDB [3] and saved as individual FASTA [37] files. Separate secondary structure predictions were run on each file using JPRED4, PSIPRED, and SPIDER2. JPRED4 predictions were run remotely using a publically downloadable scheduler available on the JPRED4 website [19]. PSIPRED and SPIDER2 calculations were run locally using the nr database [22]. Secondary structure predictions from jnetpred (JPRED), .horiz files (PSIPRED), and .spd3 files (SPIDER2) were converted into FASTA format. Each residue was assigned one of three secondary structures: ‘H’ for helix, ‘E’ for extended β-strand, and ‘C’ for coil. All experimentally determined and predicted secondary structures that were neither helix nor extended were classified as coil (including β-turns), except for chain breaks, which were annotated ‘-‘. The maximum allowable sequence length for JPRED predictions is 800 residues. Sequences that exceeded this length were pruned before being submitted to JPRED only; pruning occurred on the N-terminus, C-terminus or both N- and C-termini depending on whether the FSR was nearer to the C-terminus, N-terminus, or middle of the protein, respectively.

### Secondary structure prediction accuracy calculations

Secondary structure prediction accuracies were calculated using the Q_total_ (or Q_3_) metric [38], in which predicted and experimentally determined secondary structures are compared one-by-one residue-by-residue. Predictions were scored as follows: (in)consistent pairwise predictions were given a score of (0)/1, summed, and normalized by the length of the sequences compared. Chain breaks were excluded from both scoring and normalization. All sequences composed of ≥10% chain breaks or more were excluded from calculations.

### Secondary structure prediction accuracy distributions

All prediction accuracy distributions were calculated on protein sequences using a sliding window in steps of one. Window size equaled the length of the FSR, as defined by [10], unless it fell below 40 residues, the minimum length required for secondary structure predictions. FSR lengths below this minimum were padded symmetrically or as symmetrically as possible if located near a terminus. All protein regions containing ≥50% of the fold-switching region were considered FSR; otherwise, they were considered NFSR. FSRs were determined using the pairwise2.align.localxs function from Biopython [39] with gap-forming score of −1 and gap-elongation score of −0.5. To generate **Fig. 2**, these calculations were run exactly as before, excluding the member of each fold-switch pair with the lower Q3. All distributions were plotted using Matplotlib [40].

### Randomly-generated secondary structure predictions

First, we used the procedure described in *Secondary structure predictions of whole fold-switching proteins* to predict the secondary structures of 226 proteins with high likelihood of not switching folds [10]. Segments were selected from 10 random regions of each protein. Segment lengths were randomly selected from the distribution of FSR lengths from the 192 proteins described previously. Secondary structure prediction accuracies were calculated using the Q3 metric [38].

### Kolmogorov-Smirnov (KS) statistics and PDB Bias

We found the KS test to give implausibly low p-values for large distributions. To minimize this effect, we used the size of the smaller distribution twice, instead of using the sizes of the smaller and larger distributions once each.

To determine PDB bias, we BLASTed the sequences of our 192 proteins against the PDB and compared the structures of all hits with the structures of both fold-switch-pair conformations. Hits were grouped with the fold-switch-pair conformation to which they had the highest secondary structure similarity [10].

### Secondary structure distributions of isolated FSRs and NFSRs

Sequences of FSR fragments, padded to 40 residues when necessary (*Secondary structure accuracy distributions*), were excised from their parent sequences and saved as individual FASTA files. NFSR fragments were generated by excising fragments from 10 random locations in each non-fold-switching protein, each saved to an individual FASTA file. Fragment lengths were randomly selected from the FSR length distribution described previously. Secondary structure predictions were run on each FASTA file as described previously. N/FSR distributions were generated by determining the Q_3_ score between the secondary structure predictions of isolated N/FSR fragments and N/FSR fragments in context. Only FSRs with prediction accuracies <0.8 for both fold-switch-pair conformations were included in the distributions.

